# Early life exposure to high fructose diet induces metabolic dysregulation associated with sex-specific cognitive impairment in adolescent rats

**DOI:** 10.1101/2021.07.20.453114

**Authors:** Catherine E Barrett, Megan Jiang, Brendan G O’Flaherty, Brian Dias, Donald G Rainnie, Larry J Young, Aurelie Menigoz

## Abstract

**Background:** The incidence of adolescent mental health disorders is on the rise. Epidemiological studies suggest that poor nutrition is a significant contributor to this public health crisis, specifically through exposure to high level of dietary sugar, including fructose, during critical periods of development. Previous studies have shown that elevated fructose exposure during adolescence disrupts mental health. Further, it seems that infants display the highest level of exposure to fructose based on nutritional surveys. Despite these data, it is currently unknown how fructose exposure, specifically during infancy, may impact adolescent mental health.

**Methods:** We developed an experimental protocol in rats to investigate the effects of fructose exposure during infancy on behavioral, cognitive and metabolic endpoints in adolescence. Specifically, rat pups were exposed to fructose from birth until weaning through maternal diet. Metabolic assays, quantitative PCR and behavioral protocols such as open field, elevated O maze and a Go/ No-Go operant task, were used to determine whether high fructose exposure during infancy may set the stage for behavioral and metabolic dysfunction in adolescence.

**Results:** We found that exposing rats to high fructose from birth to weaning resulted in higher circulating glucose, insulin and leptin levels in adolescence. High fructose during infancy also increased bodyweight, disrupted metabolic homeostasis in the basolateral amygdala (BLA) as indicated by decreased activity of the cellular energy sensor AMPK, and impaired attention and impulsivity in a male-specific manner. This impaired attention observed in adolescent male rats following neonatal fructose exposure was partially rescued by viral-mediated, *in vivo* expression of a constitutively active form of AMPK in principal neurons of the BLA.

**Conclusion:** Our results suggest that exposure to high level of fructose during infancy may impact adolescent mental health in a male-specific manner and that manipulation of AMPK activity may mitigate this impact.

## INTRODUCTION

One in four adolescents living in the USA suffers from a diagnosable mental health disorder [1–3]. Mental health disorders in adolescence can have a profound impact on the individual, reduced educational achievement, substance abuse problems and an increased risk for poor physical health [3, 4]. Growing epidemiological and preclinical evidence support the association between poor nutrition and detrimental health conditions such as asthma or cardiovascular and metabolic disorders [5–7]. Unhealthy dietary choices during early life may also negatively impact brain function and disrupt behavior [8–14].

Over the last 5 decades, with the increased use of high fructose corn syrup as a sweetener in food and beverages, the daily dietary fructose intake in the US population has increased by more than 50%, and excessive fructose intake has become the most common nutritional insult experienced by children [15, 16]. Current epidemiological data have demonstrated the adverse association between high level of fructose consumption and disruption of cognitive and emotional behavior [8, 10, 17, 18]. Studies in rodents suggest that high levels of fructose during adolescence or adulthood impact emotional and cognitive behavior, in part through disruption of neuroendocrine pathways and alteration of neuronal activity and gene expression [11-13, 19]. In contrast, less is known about how fructose exposure during infancy can impact long-term behavior in adolescence. This gap in our knowledge deserves significant attention because infants and toddlers have the highest levels of fructose consumption when normalized by body weight, via exposure through breastmilk, commercial formula or processed baby food” [15, 20-22]. Nutritional surveys indicate that infants and toddlers consume on average 90g of fructose per day, which provide 30% of their energy intake, more than 3 times the recommended intake for children 2 years and older. Moreover, recent research suggests that fructose consumption may contribute to health disparities, as the dietary fructose intake vary widely among ethnic and racial groups, with Non-Hispanic Black infants consuming the most [23]. During the first 1000 days of life, the infant brain is highly plastic and develops rapidly which is associated with a very high metabolic demand. Consequently, early unfavorable conditions such as nutritional insults can impair brain organization during a critical period for cognitive health development [24]. These alarming statistics make it imperative for us to understand how infant fructose exposure may negatively impact mental health in adolescence.

We developed an experimental protocol in which rats were exposed to fructose during infancy, via maternal diet. We hypothesized that fructose exposure from birth through weaning would disrupt metabolism and alter cognitive and affective behavior in adolescent rats. To test this hypothesis, we followed the metabolic, behavioral and physiological sequelae of fructose exposure during infancy into adolescence, measuring body weight, glucose, insulin, and leptin levels and quantifying locomotor activity, social behavior, attention and impulsivity. Finally, with a focus on the basolateral amygdala (BLA), a brain region involved in the pathophysiology of mental health disorders, we examined whether the metabolic homeostasis of neurons was affected by fructose exposure by measuring the activity of the ubiquitous sensor of metabolic energy: AMP activated protein kinase (AMPK). Moreover, we tested whether viral vector-based manipulation of AMPK activity would mitigate the impact of early life fructose exposure on behavior in adolescence.

## METHODS

### Animal husbandry and diet manipulation

In this model of early life exposure to fructose, Sprague Dawley rat dams were fed a diet of control chow or 55%Cal high fructose chow (HFrD) (Research diets D05111802). The dams were fed the experimental or control diet from post-natal day 1 (P1) until weaning of their pups (P21). All experimental protocols conformed to the National Institutes of Health Guidelines for the Care and Use of Laboratory Animals and were approved by the Institutional Animal Care and Use Committee of Emory University. For details on methods, see Supplementary Materials.

### Experimental design

Animals were exposed to HFrD or control chow from P1 to P21. Litters were weaned at P21 and placed on control chow regardless of their diet during lactation. Metabolism was assessed at the end of infancy (P18) and the beginning of adolescence (P42). Behavior was assessed from P42 till P63. (Fig. 1).

**Figure 1:**
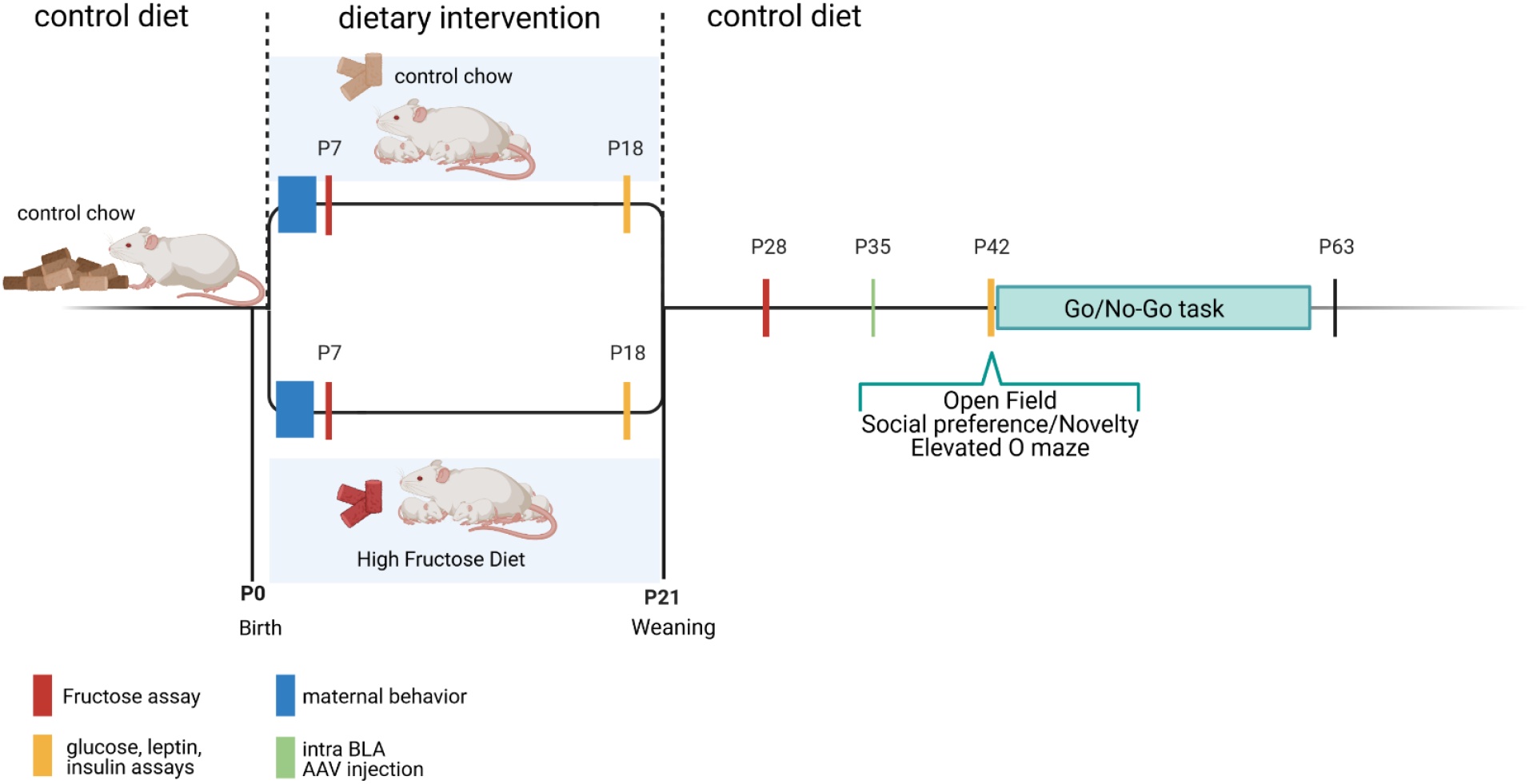
Timeline of experimental designs. Litters from a same cohort were used across multiple experimental designs to avoid cohort effect. HFrD = High Fructose Diet; PND = post-natal day.

### Metabolic assessment

Blood glucose was assessed by tail prick using a Freestyle glucometer (Abbott, IL). Blood fructose levels were measured using an enzymatic assay (EnzyChrom™ Fructose Assay Kit, BioAssay systems, Hayward, CA). Circulating plasma insulin and leptin levels were measured via ELISA (CrystalChem, Downers Grove, IL) according to the manufacturer’s instructions.

### Behavior

Starting at P40 (adolescence) locomotion, and exploratory behavior (open field test), social behavior (social preference, social novelty) and anxiety-like and risk-taking behavior (elevated O maze), were assessed over a period of 2 days. Activity within the mazes was video-recorded and analyzed off-line using TopScan (CleverSys, Reston, VA.). Learning, attention and impulsivity behavior (Go/No-Go task) were assessed from P42 to P63 using an operant conditioning chamber (Med Associates Inc). Detailed description for each test is available in Supplementary Materials.

### Stereotaxic surgery

For AMPK activity manipulation, an AAV (adeno-associated virus) vector expressing a constitutively active form of AMPKa (AMPKα2_1-312_) and GFP (Green Fluorescent Protein) or a control AAV vector (Addgene: plasmid # 60127 and # 50465) were injected bilaterally in the BLA at P35. Animals were allowed one week for recovery and sufficient time for optimal expression of the viral vector prior to the start of the behavior assessment at P42. Upon completion of the operant conditioning task, injection sites were verified as previously described by our group [25].

### Single-cell quantitative PCR

PCR was performed as previously described [26]. Cytoplasm was collected by a patch pipette under visual guidance by applying light suction. RNA was reverse transcribed into cDNA and preamplified using the Single Cell to CT kit (Life Technologies, CA, USA). Relative expression levels of AMPK subunits were determined by real-time PCR, using Universal TaqMan MasterMix (2x concentrated, Life Technologies) and TaqMan assay (20x concentrated, Life Technologies) using the ΔCt method [27, 28].

### Western blot

Total protein was extracted from tissue punched frozen BLA sections to determine the expression of Acetyl-CoA Carboxylase (ACC) and phospho-ACC. Westerns blots were performed and normalized against b-actin (1:10,000, MABT825, Millipore, Billerica, MA, USA) as previously reported by our group [25]. The relative integrated intensity value (IIV) for each sample was measured using the Alpha Innotech Fluorochem imaging system.

### Statistical analyses

Statistical analyses were carried out using Prism 8 (GraphPad Software Inc., San Diego, CA.). In this study, sample size is the number of litters. Three pups of each sex per litter were tested per experiment and for statistical purposes were considered as subsamples [29]. Two-way ANOVA, two-way repeated measure ANOVA and three-way ANOVA with Sidak’s post hoc analyses were performed as needed to analyze main effects of sex, diet, time or trials on the measured or recorded outcomes. An alpha level of 0.05 was used for all statistical tests for behavioral and the standard deviation of the mean (SD) was reported for the error.

## RESULTS

### Early life exposure to high fructose diet alters metabolism in infancy and adolescence

We did not observe any difference in body weight gain between fructose exposed and control rats during lactation. In contrast, post-weaning body weight gain analyses displayed an interaction of sex and diet ((F_9,194_ = 57.96; *p* < 0.0001). Sidak’s *post hoc* revealed that this interaction was driven by the HFrD males gaining weight at a significantly higher rate (*p*= <0.0001) than the control group; starting at P48 (Fig. 2a), indicating physiological consequences of prior fructose exposure. No differences were detected among the females (F_1,22_= 1.176; *p* = 0.2899). Interestingly, two-way ANOVA indicated that early life exposure to fructose significantly increased the fasting plasma fructose, glucose and insulin levels during infancy, regardless of sex. However, these increases did not persist into adolescence (fructose: F_3,22_= 22.24 *p*<0.001, P18 *p* < 0.0001, P42 *p* = 0.1585; glucose: F_3,74_ = 8.894, p < 0.0001, P18: *p*<0.001, P42: *p* =0.519; insulin: F_3,44_ = 28.76, p <0.0001, P18: *p*<0.0001, P42: *p*=0.9993) (Fig. 2b,c,d). In contrast, plasma leptin levels did not differ during lactation, however, it was significantly augmented in adolescent HFrD animals, regardless of sex (F_3,85_ = 20.45, p <0.0001; P18: *p* =0.9252, P42: *p*< 0.0001; Fig. 2e).

**Figure 2:**
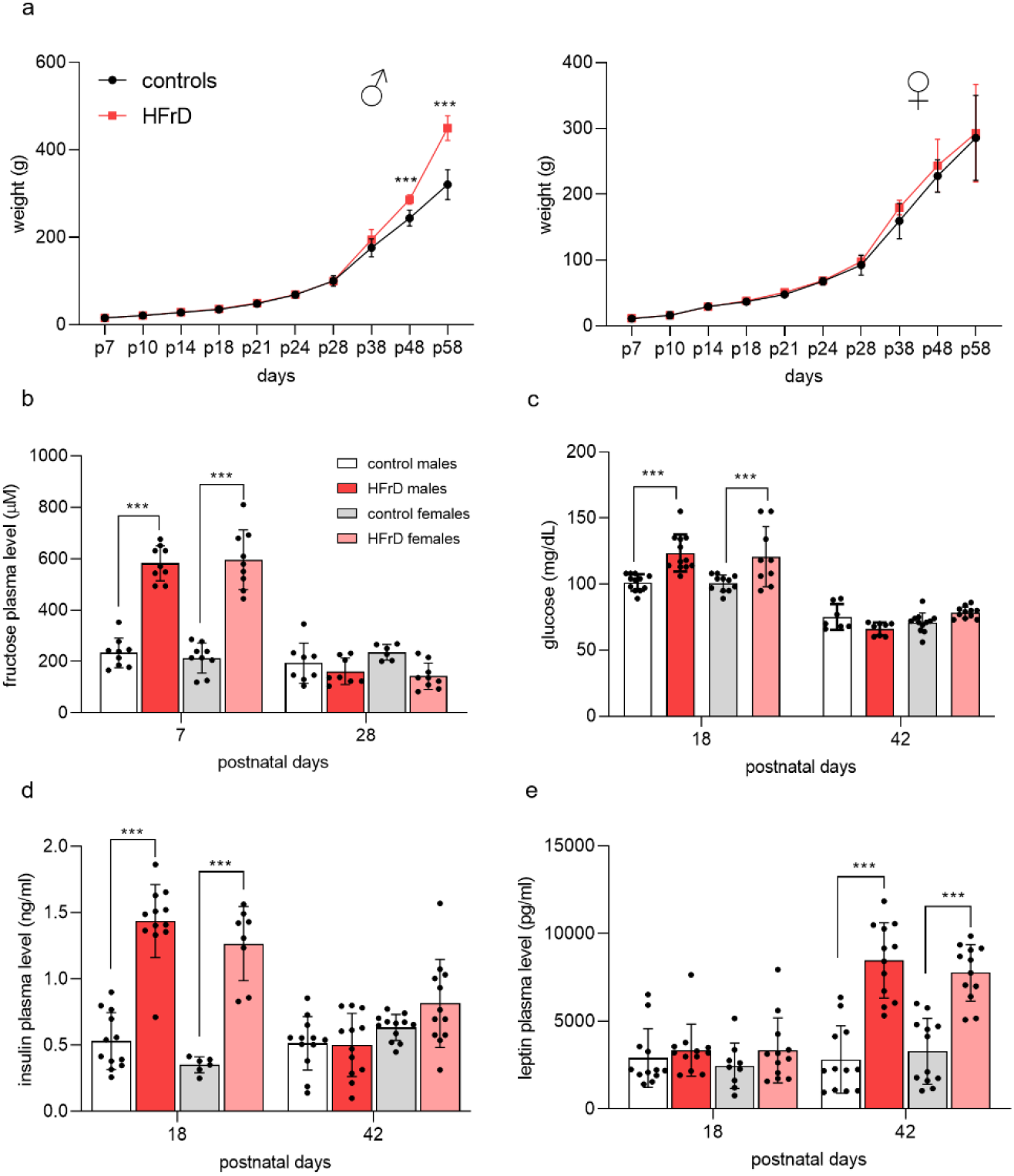
Offspring metabolic state after early life exposure to high fructose diet (HFrD). Offspring body weights graphs from birth to 58 days of age display weight gain for males and females from controls (•) and HFrD (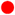) groups, with HFrD males showing an accelerated growth starting at adolescence (2-way ANOVA) (**a**). Histograms of circulating plasma levels of fructose, glucose, insulin and leptin (**b-e**) display effect of HFrD on glucose homeostasis, with HFrD groups having significantly higher glycemia and insulin and leptin levels (2-way ANOVA with Sidak’s post hoc). Values are mean ± SD of n = 12 litters/group. Asterisks indicate significant comparisons between HFrD and control groups ***p ≤ 0.001.

### Early life exposure to HFrD alters locomotion without affecting social behavior

To control for diet-induced changes in maternal care we next compared nesting and weaning behavior between the two groups. No significant difference was detected in frequency of dam on nest, arch back nursing, or passive nursing (Fig. 3a). We next examined the effect of HFrD exposure on anxiety-like and locomotor behaviors during adolescence (P42-45) using the open field, elevated O maze and social novelty or preference test. Three-way ANOVA analyses of open field activity revealed no difference on thigmotaxis but a significant main effect of diet on the total distance traveled (m) (con: 15.11±0.87; HFrD: 24.62±1.1; F_1,31_=60.87;*p*<0.0001) (Fig. 3b,c). HFrD exposed animals spent significantly more time in the open arms of the elevated O maze compared to controls regardless of sex (con: 166.3±3.239; HFrD: 215.4±1.12; F_1,31_ = 32.82; p < 0.0001), suggesting elevated exploratory behavior (Fig. 3d). Finally, diet and sex did not impact the social or novelty preference behavior of the adolescent animals (diet: F_1,68_ = 0.0125; *p* = 0.9111; sex: *F*_1,50_ = 1.67; *p* = 0.2107) (Fig. 3e).

**Figure 3:**
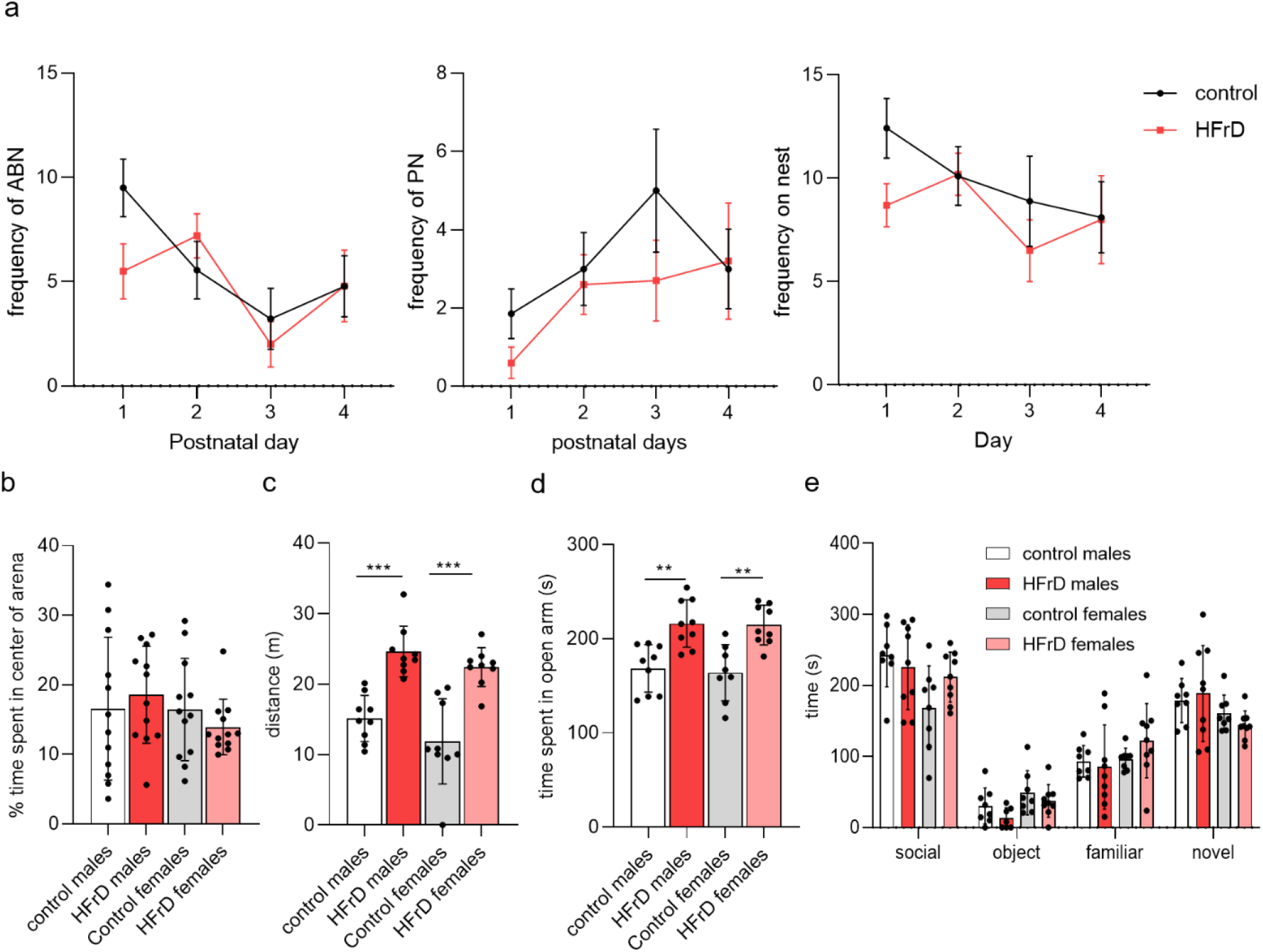
Behavioral phenotype of early life exposed HFrD animals. Frequencies of arch-back nursing (ABN), passive nursing (PN), and on the nest were unaltered between control (•) and fructose (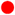) fed dams (**a**). Adolescent HFrD males (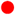) andfemales (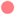) showed no difference in thigmotaxis compared to control fed males (•) and females (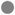) (**b**), however the locomotion of both HFrD groups was significantly higher than the control groups (two-way anova with Sidak’s post hoc (**c**)). HFrD males and females spent more time in the open portion of the elevated O maze (EOM) (one-way anova with Sidak’s post hoc (**d**). All groups showed preference for a social stimulus in the social preference test and no differences were detected in latency to approach or time spent in proximity to a novel conspecific during the social novelty component (**e**). Values are mean ± SD of n = 12 litters/group. Asterisks indicate significant comparisons between HFrD and control groups *p < 0.05; **p < 0.01; ***p ≤ 0.001

### HFrD decreases attention and inhibition in a sex specific manner

We next assessed the effect of fructose exposure on learning, attention and impulsive behavior in adolescent rats using a Go-No-Go paradigm (Fig. 4a). In phase 1, HFrD exposure had no significant effect on the ability of the animals to learn the operant association between lever pressing and reward delivery. All groups reached criterion in a similar time frame (Log-Rank chi-square = 0.5784, df = 3, p = 0.9014; Fig. 4b) and with the same accuracy (diet: F_1,22_=1.43, *p*=0.2433; sex: F1,_22_=0.3967, *p*=0.3967; Fig. 4c). Strikingly, 2-way ANOVA showed a significant interaction of diet and sex for the time to reach completion criterion (F_1,84_=10.85, *p*=0.0014; Fig. 4b), the accuracy(F F_1,22_= 7.244, *p*= 0.0133; Fig. 4c) and the error rate (F_1,31_ = 7.797, *p*=0.0089; Fig. 4d) in phase 2, suggesting an impairment in attention behavior (Fig.4b, d). *Post-hoc* analyses revealed an increase both in the time to reach criterion and in the error rate (*p*= 0.0104 and *p*= 0.0480, respectively) in HFrD exposed males. In phase 3, a stop auditory cue was introduced to examine the effect of diet and sex on impulsive behavior. Two-way ANOVA and post-hoc analyses revealed a significant diet by sex interaction on the success rate in No-Go trials (F_1,22_=11.22, *p*=0.0029) driven by worse performance in the No-Go trials of the HFrD males compared to the other groups (Fig.4e), suggesting a higher level of impulsivity. No difference was detected in the Go trial in phase 3. Analysis of the omission rate and average response time across the 3 phases (Table 1) indicated no difference in motivation throughout the task.

**Figure 4:**
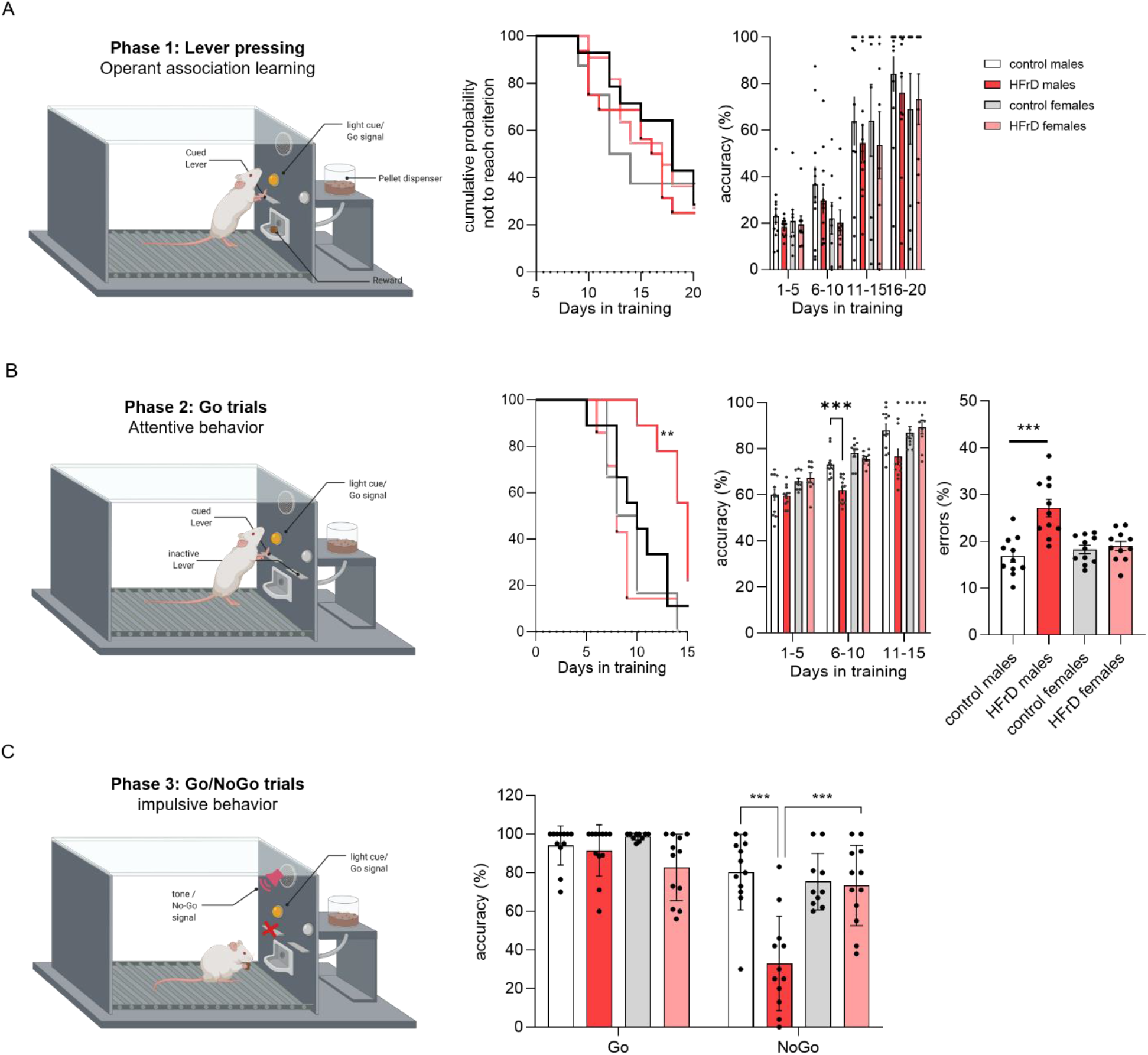
Effects of HFrD early life exposure on Go/No-Go task performance. Schematic of the Go/No-Go paradigm design. The task comprised 3 phases designed to test learning, attention and impulsivity. Learning of the operant association lever pressing + reward (left panel) was assessed Kaplan-Meier survival curves showing the cumulative probability of adolescent HFrD males (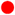) and females (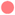) and control fed males (•) and females (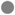) subjects not achieving criterion across training sessions (middle panel) and the corresponding accuracy data (right panel) (**A**). Attentive behavior was assessed in phase 2 through cue discrimination during go trials (left panel), multiple comparisons indicate that HFrD males took significantly longer to reach criterion, had lower accuracy of response (middle panels) and presented a significantly higher average error rate (right panel) (**B**). Impulsivity was assessed through inhibitory control with the introduction of an auditory stop signal (phase 3) (left panel). Performances in phase 3 as measured by the accuracy of the responses on Go and No-Go trials revealed that HFrD males inhibited their planned response in No-Go trials significantly less than the other groups (right panel) (**C**). Single dots represent individual litters and bars are average mean ± SD of n = 12 litters/group. Asterisks indicate significant comparisons between groups *p < 0.05; **p < 0.01; ***p ≤ 0.001.

**Table 1:** comparison of the performances in Go/No Go operant task. All data are expressed as mean ±SD.

|  | PHASE 1 |  |  | PHASE 2 |  |  | PHASE 3 |  |
| --- | --- | --- | --- | --- | --- | --- | --- | --- |
|  | sessions to reach criterion (days) | Omission (nb/session) | Response latency (ms) | sessions to reach criterion (days) | Omission (nb/session) | Response latency (ms) | Omission (nb/session) | Response latency (ms) |
| control |  |  |  |  |  |  |  |  |
| Males | 14.667±5.483 | 4.92±3.806 | 13.177±3.671 | 9.309±1.456 | 0.633±0.57 | 5.009±1.166 | 1.167±1.13 | 4.386±2.372 |
| females | 12.811±2.267 | 7.292±2.79 | 19.360±2.246 | 10.420±2.427 | 0.451±0.653 | 3.169±0.531 | 0.818±0.367 | 2.873±2.806 |
| HFrD |  |  |  |  |  |  |  |  |
| Males | 15.250±5.259 | 4.724±4.259 | 12.225±4.612 | 14.248±2.368** | 0.51±0.466 | 4.611±0.955 | 0.813±0.948 | 4.16±1.716 |
| Females | 15.008±3.163 | 9.132±1.723 | 17.251±3.937 | 10.318±2.786 | 1.018±1.1 | 3.967±0.613 | 1.075±0.782 | 4.977±3.821 |
| Control males + |  |  |  |  |  |  |  |  |
| eGFP | 15.833 ±2.137 | 7.377±0.685 | 11.608±1.202 | 9.194±3.583 | 0.569±0.317 | 3.262±0.481 | 0.118±0.145 | 2.851±0.73 |
| AMPK$\alpha_{1-312}$ | 16.5± 2.871 | 7.089±1.3 | 11.463±1.738 | 10.389±2.542 | 0.542±0.292 | 3.652±1.401 | 0.079±0.097 | 2.415±0.502 |
| HFrD males + |  |  |  |  |  |  |  |  |
| eGFP | 14.667±1.935 | 8.075±0.554 | 10.606±1.548 | 14.777±1.292 | 0.889±0.574 | 4.387±0.549 | 0.382±0.461 | 3.365±0.372 |
| AMPK$\alpha_{1-312}$ | 13.371±2.337 | 7.17±0.647 | 10.708±0.916 | 10.182 ± 2.16 | 0.896±1.05 | 4.638±0.799 | 0.417±0.376 | 3.07±0.928 |

### Increased AMPK activity in BLA neurons partially rescues HFrD induced attention deficit in males

Previous studies have reported that the BLA plays a key role in attentional and representational processing [30–32]. Hence, we investigated the effect of HFrD on the expression and activity of the energy sensor AMPK in BLA principal neurons [33]. At P18, all three AMPK subunits mRNA levels were significantly lower in HFrD exposed males than the levels found in BLA neurons in controls (Fig. 5a). In addition, AMPK activity was indirectly assessed using the ratio of pACC/ACC in total protein extract isolated from BLA tissue sections determined by Western-blot analysis. At P42, HFrD exposed males showed a significantly reduced ratio of pACC/ACC compared to controls (F_1,28_= 82.60, *p*<0.001), which together with the reduced AMPK subunit mRNA levels strongly suggested that HFrD exposure induced a prolonged decrease in BLA neuronal AMPK activity in males (Fig. 5b). To explore the possibility that decreased AMPK activity in BLA principal neurons contributes to HFrD-induced behavioral impairments in adolescence, we examined the effects of selectively modulating BLA neuronal AMPK activity using a AAV vector encoding a constitutively active form of AMPKα (AMPKα_1-312_) in control and HFrD males. We observed no significant effects of AMPKα_1-312_ treatment on the ability to learn the operant association (F_1, 20_ = 1.013, *p*=0.3262) in phase 1 (Fig. 5c,d). In phase 2, AMPKα_1-312_ treatment decreased the latency to reach criterion (F_1, 20_= 7.833; *p*=0.0111) and the error rate (diet: F_1, 20_= 17.79, *p*=0.0004; Diet x Treatment: F_1, 20_= 7.739, *p*=0.0115) and increased accuracy (F_1.359, 13.59_ = 4.622; *p*= 0.0407) in the HFrD males to the equivalent level of the control groups (Fig. 5c-e). Finally, three-way ANOVAs found no significant effect of treatment in the Go and No-Go trials in phase 3 (F_1, 10_ = 3.178, *p*=0.1050; Fig.5f). HFrD males, regardless of treatment, had a lower accuracy in the No-Go trials compared to the control groups (*p*= 0.0183). There was no difference in the omission rate and average response time across all the phases (Table 1).

**Figure 5:**
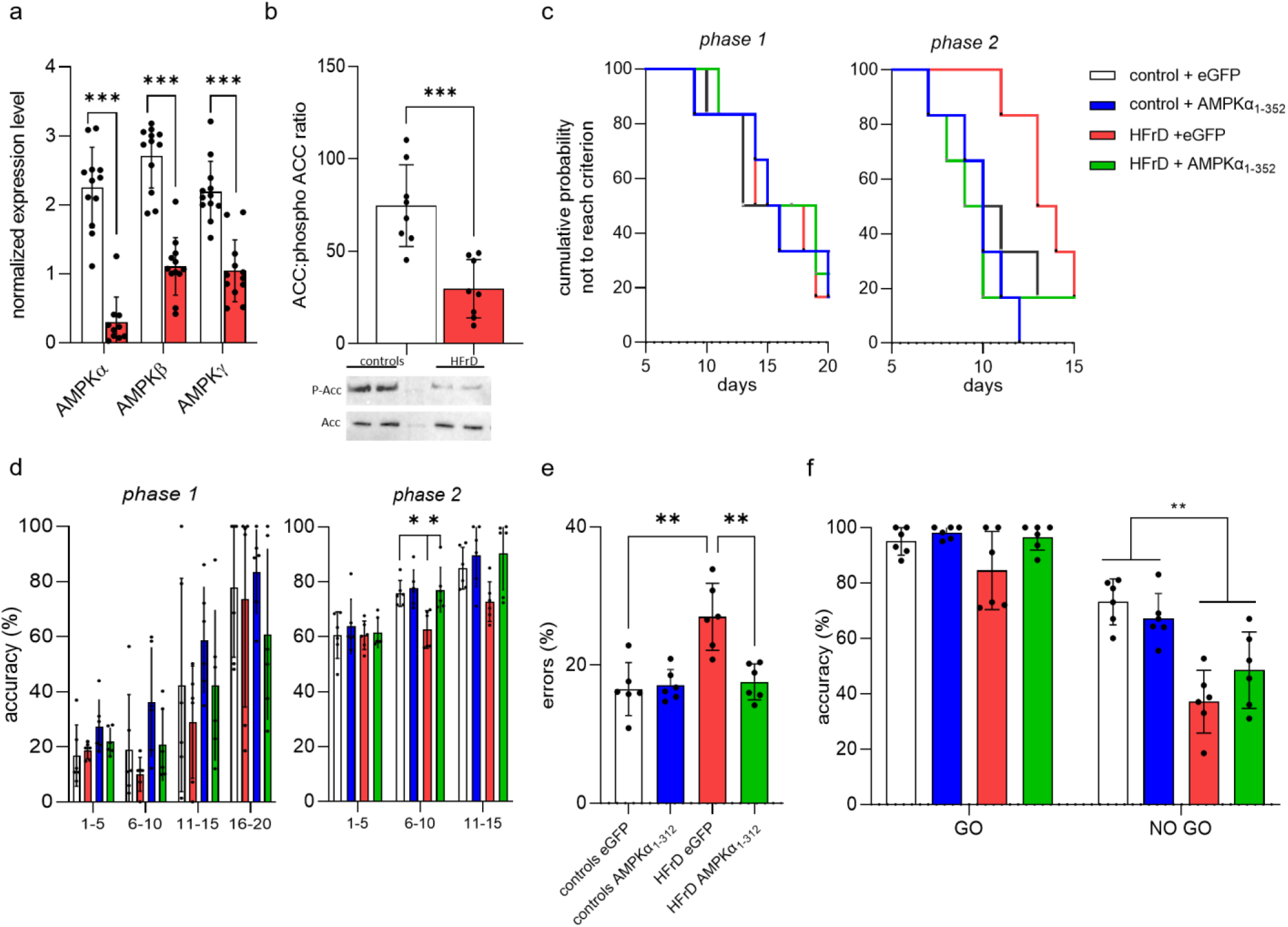
Early life HFrD exposure decreases offspring BLA AMPK expression and upregulation of its activity partially rescues diet induced behavior impairment. Normalized mRNA level (mean ± SD) of AMPK subunits from BLA principal neurons (n = 12 litter per group (3 male/litter) *** p < 0.001 vs control) (**a**) and AMPK kinase activity as measured by the ratio of phospho-Acetyl-CoA Carboxylase (ACC) determined by immunoblotting (top panel, **b**) and representative blot (bottom panel, **b**) (n = 8 litters/group (3 male/litter) *** p < 0.001 vs control). Student t-tests indicate that HFrD decreased both the mRNA expression and the activity of AMPK. Kaplan-Meier survival curves showing the cumulative probability of not reaching criterion of adolescent HFrD adolescent males treated with eGFP or AMPKα_1-312_ encoding AAV (HFrD eGFP, 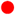 and HFrD AMPKα_1-312_, 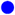 respectively) and control counterparts (control eGFP •; control AMPKα_1-312_ 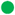) for phase 1 (left panel) and phase 2 (right panel) (**c**). Multiple comparisons indicate that HFrD AMPKα_1-312_ reached criterion in phase 2 significantly faster than HFrD eGFP. The corresponding accuracy data are shown for phase 1 and phase 2 in (**d**) left and right panels respectively. The error rate during phase 2 training revealed a significant improvement in HFrD AMPKα_1-312_ compared with HFrD eGFP (**e**). Performances in phase 3 as measured by the accuracy of the responses on Go and No-Go trials (**f**). Multiple comparisons revealed that AMPKα_1-312_ encoding AAV does not improved phase 3 performances in No Go trials for the HFrD fed males. Single dots represent individual litters and bars are average mean ± SD of n = 6 litters/group.

## DISCUSSION

In this study, we present a metabolic characterization and multi-behavioral phenotyping of adolescent rats previously exposed to high level of fructose during infancy. This fructose exposure led to long lasting changes in metabolic markers and these alterations were associated with male-specific changes in attention and impulsive behavior. Finally, exposure to high-fructose during infancy decreased expression and activity level of the metabolic regulator AMPK in the BLA, and viral-mediated increase in AMPK activity in fructose fed males partially rescued the deficit in attention and impulsive behavior. Hence, we have shown that exposure to HFrD selectively during infancy causes sex-specific alterations in metabolism and behavior that may, in part, be driven by the dysregulation of AMPK activity in the amygdala.

Consumption of fructose during infancy altered the physiological parameters assessed in this study in both male and female rats. Early life exposure to fructose led to increased circulating fructose levels at P7, which were normalized to control level after one week on control lab chow (P28), indicating that the exposure to fructose was indeed restricted to the lactation period. The presence of fructose in offspring’s blood indicates that the dam’s breastmilk composition was altered by the diet and despite an indirect exposure, the offspring did receive a direct nutritional effect of fructose. Early life HFrD exposure led to significant metabolic alterations during adolescence. Notably, starting at adolescence (P42), the fructose-exposed males became significantly heavier and displayed higher plasma leptin levels than controls, while no difference was observed in food intake. These findings are consistent with previous work in adult models of diet-induced metabolic perturbation, where increased circulating leptin levels leads to exacerbated weight gain by altering energy utilization [34, 35].

In the open field paradigm, early life fructose exposure did not lead to any difference in anxiety-like behavior compared to adolescent controls. Similarly, analysis of social interactions suggested that diet did not impact adolescent social behavior. However, early life HFrD increased the total distance traveled in the arena, suggesting an effect on locomotion. These results are generally in line with previous studies showing that high fat/sugar diet or western diet exposure occurring during the perinatal period does not diminish sociability in rodents, but could lead to hyperactive behavior [12, 36-41]. HFrD exposure increased the time spent in the open arms of the elevated O maze in adolescent rats, independent of their sex. While elevated level of novelty seeking is a behavior expected during adolescence [42], these data showed a significant increase in exploratory behavior following fructose exposure, compared to controls. This increase in risk taking behavior might be a direct manifestation of the overall increase in locomotion as observed in the open field test and does not necessarily relate to the exploratory behavior of the HFrD exposed animals. Nevertheless, the increased locomotion and risk taking behavior observed following early life exposure to HFrD are consistent with previous studies in mice showing enhanced exploratory behavior in elevated maze after exposure to overfeeding or high fat/high sugar diet [39–41]. Interestingly, adolescence or adult exposure to a high caloric diet leads to the opposite results, with an increase in anxiety [12, 43] and reduced social preference and social recognition [14, 41, 44], similar to what was seen in human studies [45, 46]. Taken together, these data suggest that the impact of the nutritional insults on animal behavior is dependent on the timing of the insult, most likely due to the susceptibility of specific neural circuits in critical developmental windows such as early life or adolescence. Alternatively, the increased exploratory drive behavior observed in our study may be due to the degree of maternal attention received during the neonatal period [47, 48]. An abrupt change in diet at the beginning of the lactation period could cause a significant stress response in the dams leading to a decrease in maternal attention. Our daily 1-hour observations of the maternal behavior throughout the first week of lactation did not reveal any difference in grooming, breastfeeding position or time spent interacting with the pups between control and fructose fed dams. These data would suggest that impaired maternal care did not contribute to the behavioral changes observed in the HFrD-exposed pups.

Clinical and pre-clinical work have previously shown that exposure to sugars during adolescence or in the perinatal period impairs attention and impulsivity [12, 17, 38, 49, 50]. In the present study, the impact of fructose exposure during lactation on cognitive function in adolescence was assessed using a Go/No-Go task, a paradigm used in neuropsychology to assess pro-attentive skills and impulsive choices [51]. In contrast, two recent studies [52, 53] reported that neonatal exposure to a high caloric diet led to impaired learning and memory. HFrD exposure during infancy did not impair the ability to form the operant association. These contrasting results might be due to differences in the type of nutritional insult used, as high caloric diet or western diet contains high levels of fat versus high levels of sugar. Those reports assessed learning using the Morris water maze while we used an operant task, therefore, we cannot rule out the possibility that the HFrD exposure may also impair hippocampal dependent learning like studies with neonatal overfeeding and western diet exposure have reported. Fructose-exposed males showed decreased attention during the acquisition of the Go signal phase, as well as increased impulsive-like behavior as measured by continued lever pressing after presentation of the Stop signal. No differences were not noted in females. These data are in line with recent work in rodents and humans; epidemiological studies showed that children exposed to elevated levels of sugar before the age of 7 had poorer executive functioning during adolescence [17, 18, 54-56]. In mice, sucrose exposure during gestation and lactation led to attentional deficits and increased impulsivity in offspring [38]. While most previous studies did not assess the interaction of diet and sex, the few studies that did, reported more prominent effects of diet on behavior in males [57–59]. These sex differences in vulnerability to diet may reflect different adaptive strategies to the environment, or be the results of the neuroprotective effects of estrogen [60, 61]. Overall, despite some evidence of its importance, the mechanisms underlying moderation of early life nutrition by sex with regard to cognition remain largely unknown.

AMPK is serine/threonine kinase that is known as a master regulator of cellular metabolism. Prior research in adolescent or adults have established that nutritional insults are commonly associated with perturbation of AMPK activity [62–65]. Recent studies have suggested that metabolic insults could lead to behavioral impairment by dysregulating the activity in key limbic regions such as the BLA [66–68]. While known for its role in fear-related behavior, the BLA also plays a critical role in regulating decision making processes, executive function and impulsivity [31, 69, 70]. For these reasons, we examined how early life exposure to fructose impact the expression and activity of AMPK in the BLA. Here, we found that fructose exposure during infancy led to a decrease in AMPK activity in the BLA. Consistent with this finding, previous studies have shown that diet-induced behavioral perturbations are associated with decreased levels of activation of AMPK [57, 58, 62-66, 71].

In order to examine any cause-effect relationship between decreased AMPK activity in the BLA and alterations in adolescent behavior, we next determined if we could rescue the behavioral deficits observed in HFrD exposed males. To this end, we treated adolescent male rats with an intra BLA injection of AAV viral vector encoding either a constitutively active form of AMPKα_(1-312)_, or GFP as a control. Viral manipulation of BLA AMPK activity did not impair the ability to learn the operant association in the Go/No-Go task. Significantly, following AMPK viral manipulation, fructose exposed males displayed similar performance levels in the phase 2 compared to control fed animals. However treated HFrD males displayed higher impulsivity in phase 3. These data suggest that augmenting AMPK activity in the BLA significantly improved the attention behavior of the fructose fed animals in the Go/No-Go task, while not affecting the impulsive behavior. Hence, in line with previous work showing that behavioral perturbations were alleviated following administration of the AMPK activator AICAR, these findings point toward a role for AMPK dysregulation in the BLA in fructose-induced attention impairment. As inhibitory control and impulsivity can be processed through a diverse neuronal circuitry [72–74], our inability to rescue fructose induced impulsivity by increasing AMPK activity in the BLA was possibly due to this specific behavioral consequence of HFrD being mediated by one or several brain regions other than the BLA, such as the prefrontal and/or the orbitofrontal cortex [31].

Growing evidence suggests that sugar consumption during adulthood significantly increases baseline neural activity of the BLA in humans and rodents alike [13, 75, 76] and aberrant activity in the BLA plays a major role in behavioral impairments [68, 77-80]. Moreover, recent studies have shown that AMPK modulates membrane excitability by directly regulating the expression of membrane bound neurotransmitter receptors and ion channels [81–83]. Our data support the novel hypothesis that decreased AMPK activity in the BLA set into motion by early life fructose exposure leads to aberrant neuronal activity in BLA principal neurons, underlying the attentional deficits observed in fructose fed animals. Consequently, future studies should directly assess and manipulate neuronal activity in the BLA of adolescent rats following early life fructose exposure, while querying and rescuing deficits in attentional behavior.

To conclude, early lifestyle factors can be detrimental to neurodevelopment and predispose children to subsequent mental health disorders. Here, we provide experimental evidence for a role of infant fructose exposure on behavioral perturbations in adolescence. While previous studies have investigated the effects of a dietary disturbances during adolescence or adulthood on behavior, we focused our attention on the effect of fructose exposure during infancy on behavior during adolescence. We demonstrated that early life exposure to high fructose diet leads to attentive and impulsive behavior impairments in a sex-specific manner, in part via altering AMPK activity in the BLA. Considering the lack of guidelines regarding dietary fructose intake for infants and toddlers, how pervasive fructose is in US diet and how fructose consumption starts early in life, and its potential role in future vulnerability to mental health disorders, our data emphasize the need to study early life nutritional insults independently from insults during the adolescent period.

## Acknowledgments

We would like to thank Rachel Chernoff for technical assistance and the animal caretakers at Yerkes National Primate Research Center for excellent animal care. This research was supported by National Institutes of Health Grants R01MH0069852 to DGR and LJY, and P51 OD011132 to YNPRC. The authors declare no conflict of interest.

