## supplemental material for "Early life exposure to high fructose diet induces metabolic dysregulation associated with sex-specific cognitive impairment in adolescent rats"

**METHODS**

**Experimental subjects**

***Animal husbandry:*** All animal experiments followed the guidelines described in the NIH Guide for the Care and Use of Laboratory Animals (NIH Publications No. 8023). Animals were housed in the Yerkes National Primate Research Center vivarium at Emory University, kept on an artificial regular 12:12-h light cycle and were housed in ventilated cages kept at 22 °C. A total of 52 Sprague Dawley rat litters were used in the following experiments. All experimental protocols conformed to the National Institutes of Health Guidelines for the Care and Use of Laboratory Animals and were approved by the Institutional Animal Care and Use Committee of Emory University. For details on methods, see Supplementary Materials.

***Experimental design:*** In this model of early life exposure to fructose, the experimental animals are offspring reared by rat dams placed on a high fructose diet (HFrD) (described below) or control chow from post-natal day 1 (P1) until weaning (P21). Dams were ordered from Charles River (Wilmington, MA) and arrived on gestational day E7. At birth, half of the dams stayed on control chow while the other half was switched to high-fructose diet. The litters were randomly assigned to either diet to balance litter size across groups. Maternal behavior was assessed from P2 to P5 to ensure that the dietary perturbation did not alter maternal care. At P21, the litters were weaned, placed on control chow regardless of their group assignment and housed by 3 with same sex litter mates. When the litter size allowed for it (rat litter sizes varied between 10 and 14 pups), pups from a litter were used across multiple experiments. Six independent cohorts of litters were used for this study. The metabolic assessments were measured at end of infancy (P18), and beginning of adolescence (P42), except for fructose blood levels which were assessed at P7 and at P28, i.e one week after weaning animals to control diet. Behavior was assessed from P42 till P63 and was preceded by 5 days of handling to habituate the animals (Fig. 1).

**Diet and metabolic measurement**

***High Fructose Diet:*** The fructose diet used (Research diets D05111802) was 55% fructose while the 'control' chow (Lab Diet 5001) contained only 0.30% fructose. Previously, this fructose diet has been shown to reliably elicit metabolic changes (hyperglycemia and insulin resistance) in rats [13]. Both diets contain comparable levels of vitamins and minerals necessary to maintain rodent health and were reviewed by veterinary staff and approved by IACUC. The details of the macronutrients of each diet have been listed in Supplemental Table 1. Except when necessary for blood glucose measurement, animals had *ad libitum* access to food and water.

***Metabolic assessment:*** Blood glucose was tested weekly after a 6-hours fast by tail prick using a Freestyle glucometer. Animal weights were taken concurrently with glucose readings. Blood fructose levels were measured at P7 and P28 from plasma collected between 8 and 10am using an enzymatic assay (EnzyChrom™ Fructose Assay Kit, BioAssay systems, Hayward, CA). Insulin and leptin were measured in plasma collected between 8 and 10 am, via ELISA (CrystalChem, Downers Grove, IL) according to the manufacturers instructions. All samples were run in triplicate and any sample with intra-assay coefficient of variation > 15% was excluded from the analysis.

**Behavior**

***Maternal behavior:*** Home cage observations were conducted daily from P1 to P4 (17:00–18:00) on control and fructose fed dams. Position of dams on the litter (on or off), nursing style (arched-back, blanket, passive), and licking/grooming pups were made every 4 min (15 obs/hour).

***Open field and social interaction:*** At P40, rats were placed in the center of the testing arena (27 × 90 × 90cm) for 10 min under red light illumination, and the percentage of time spent in the center of the field (middle 50%) was used as a measure of anxiety. That time also served as habituation to the novel environment prior to social testing. Subsequently, a cage containing a same-sex, same-age conspecific and an empty cage were placed at opposite corners of the arena to test the subject animal's preference to spend time in proximity to another rat or an object. After 5 min, a novel same-sex, same-age conspecific was put into the empty cage, and the preference for a novel or familiar animal was assessed over 5 min. For analysis, the arena was divided into four quadrants, each approximately 45 × 45 cm. The social preference portion contained a social zone, an object zone, and two empty zones. Subsequently, the social novel portion contained a familiar zone, a novel zone, and two empty zones. Durations in, entry bouts and distances walked into the zones were analyzed using the automated system TopScan (Cleversys, Reston, VA).

***Elevated O Maze (EOM):*** The EOM is a variant of the elevated plus maze consisting of a circular platform (50 cm diameter with platform 5 cm wide) elevated 50 cm above the floor with two opposing open (no walls) and enclosed (30-cm high walls) quadrants (Stoetling, IL, USA). Rats were placed in the center of one of the open regions of the EOM and allowed to explore for 5 minutes. The time spent in the open quadrants of the EOM was used as a measure of anxiety. All activity within the maze was recorded and analyzed off-line using TopScan (CleverSys, Reston, VA.).

***Operant condition: Go – No-Go:*** Rats began training for the operant conditioning task Go-No-Go at P42, using illuminated Med-Associates operant conditioning chambers ((55.69 x 38.1 x 35.56 cm) (Med Associates Inc) equipped with a nose poke recess, 2 retractable levers and cue lights, a speaker for tone delivery and a modular pellet dispenser for food reward delivery. Training sessions happened following this schedule: two sessions per day, 5 days a week. Sessions ended when the maximum of 50 successful trials was reached or after 20 min. A trial was defined as the period between the house-light on (start of trial) and the house-light off (end of trial). The trial duration depended on the performance but never exceeded 1 min. The task was divided in 3 phases: Lever Pressing learning, Go Signal and No-Go signal. Performance criterion to move from one phase to the next are described below.

In phase 1, the animals were trained to nose poke the magazine at the beginning of the trial ( signaled by the houselight) to trigger the lever extension (one lever cued with a light signal, presented left or right in a pseudorandom fashion) and lever press to trigger reward (chocolate flavored food pellet) delivery. A correct lever press delivered a reward (1 pellet) on a FR1 ratio, with an inter-trial interval set at 10 sec. Failure to nose poke or lever press was recorded as ‘omission’ and triggered a 10 sec time-out. This phase allowed the animals to habituate to the chamber and learn the operant association between lever pressing and the reward delivery. Accuracy is defined as the percentage of reward obtained versus the number of triggered trials. The animals moved to the next phase once they reached 80% accuracy for 3 consecutive sessions. In phase 2, nose poke triggered extension of both levers (only one active cued lever, left or right in a pseudorandom fashion). Pressing cued lever led to reward delivery. Non-active lever pressing ended the trial, added a 10sec time out and was recorded as ‘error’. Failing to trigger the trial or lever press was recorded as ‘omission’. The error rate was calculated as follows:

Errors/(Rewards+Errors). In phase 3, nose poke triggered extension of one lever. Here, 70% of the trials were Go trials similar to phase 2. The remaining 30% of the trials were No-Go where lever extension was followed by the presentation of a stop signal indicating the need to inhibit lever pressing. The stop signal consisted of a tone delivered after a delay established for each animal to be 0.5sec under their reaction time in phase 2. Lever pressing during a No-Go trial was recorded as 'false alarm'. Each session in phase 3 started with 2 forced No-Go trials to ensure the animals were exposed to the consequence of the stop signal. The animals are trained for 4 consecutive days and day 5 was used to test their accuracy in the Go and No-Go trials.

**Viral injection**

At P35, 500nl of AAV-CaMKIIa-AMPKa2(1-312)-GFP (constitutively active form of AMPKa) or AAV-CaMKIIa-mCherry (control) were infused (bilaterally in BLA (AP:- 3.6 mm, ML:+ 3.83 mm, DV: - 9.45 mm, 6° coronal angle) using stereotaxic surgery. All coordinates were measured from the surface of bregma. Metacam (2 mg/kg, *i.p.*, Metacam, Boehringer Ingelheim Vetmedica, Inc.) was administered daily for 2 days post-surgery. The animals were allowed one week for recovery and sufficient time for optimal expression of the viral vector. To avoid litter effect, every litter received both treatments, however, to account for litter size variability, only 2 males per litter were injected with either virus. Operant conditioning started at P42 as previously described. Upon completion of the operant conditioning task, following the standard 4% PFA fixation protocol the brains were removed, post-fixed, and sliced to verify viral injection site as previously described by our group [24].

**Perfusion, sectioning and immunohistochemistry**

Rats were euthanized via pentobarbital (100 mg/kg, SomnaSol, Henry Schein Animal Health) and perfused with phosphate buffered saline (PBS), followed by 4% paraformaldehyde (pH 7.4 in PBS, Sigma-Aldrich). Brains were removed and stored at 4°C in 4% paraformaldehyde overnight and transferred to 30% sucrose solution until they reached an isosmotic state. Brains were sectioned into 50 µm slices using a freezing sliding microtome (Microm). For staining, sections were blocked and permeabilized with 3% BSA and 0.2% Triton X-100, then incubated overnight at 4 °C with appropriate primary antibodies (rabbit anti-GFP, 1:1000, A11122, Life Technologies Carlsbad, CA). The following day, sections were washed and incubated with Alexa 488 goat anti-rabbit (Life Technologies) for 2 h at room temperature. Slices were mounted on gelatin coated microscope slides and coverslipped. High-resolution images were obtained using confocal spinning disk laser microscopy and captured using an Orca R2 cooled CCD camera (Hamamatsu, Bridgewater, NJ) mounted on a Leica DM5500B microscope (Leica Microsystems, Bannockburn, IL) equipped with a CSU10B Spinning Disk (Yokagawa Electronic Corporation, Tokyo, Japan).

**Single-cell quantitative PCR**

This was performed following the protocol previously established by our group [25]. Cytoplasm was collected by a patch pipette under visual guidance by applying light suction until the cell had visibly shrunken. The content of the pipette was processed using the Single cell to CT kit (Life Technologies, CA, USA) to produce cDNA and amplified using Preamplification Master Mix (Thermofisher, CA, USA). Relative expression levels of AMPK subunits were determined by real-time PCR, using Universal TaqMan MasterMix (2x concentrated, Life Technologies) and TaqMan assay (20x concentrated, Life Technologies). The reactions were performed with a 7500 Fast Real-Time PCR System (Life Technologies). GAPDH and B2M, selected using the geNorm application, were used as endogenous controls for calibration. The relative expression of mRNA was normalized to the geometric mean of the calibrators, using the ΔCt method (Livak and Schmittgen, 2001; Vandesompele et al., 2002). Sample sizes ranged from 47 – 52 cells per group. The comparative analysis was done using geometric mean rather than the arithmetic mean to be consistent with the lognormal distribution of the level of expression. Preamplification uniformity was assessed using the ΔΔCt method. Relative expression levels of AMPK subunits were determined by real-time PCR, using Universal TaqMan MasterMix (2x concentrated, Life Technologies) and TaqMan assay (20x concentrated, Life Technologies).

**Western blot**

Total protein was extracted from tissue punched frozen BLA sections to determine the expression of Acetyl-CoA Carboxylase (ACC) and phospho-ACC. The phosphorylation of ACC has been shown to strongly correlate with changes in AMPK activity (Park SH, 1985) and was therefore used as a proxy for AMPK activity. Westerns blots were performed and normalized against b-actin (1:10,000, MABT825, Millipore, Billerica, MA, USA) as previously reported by our group [24]. For detection of pACC, NaF (50 mM), an effective protein phosphatase inhibitor, was added to all buffers. ACC and Phospho-ACC antibodies were obtained from Cell Signaling, Inc

(Cell Signaling, MA, USA): ACC (1:500, #3662); Phospho-ACC (1:500, #11818). The relative integrated intensity value (IIV) for each sample was analyzed using the Alpha Innotech Fluorochem imaging system.

**Statistical analyses**

Statistical analyses were carried out using Prism 8 (GraphPad Software Inc., San Diego, CA.). In this study, sample size is the number of litters and 3 pups of each sex per litter were tested per experiment and for statistical purposes are considered as subsamples [26]. Two-way ANOVA, two-way repeated measure ANOVA and three-way ANOVA with Sidak's *post hoc* analyses were performed as needed to analyze main effects of sex, diet, time or trials on the measured or recorded outcomes. An alpha level of 0.05 was used for all statistical tests for behavioral and the standard deviation of the mean (SD) was reported for the error.

**Supplementary table 1:** Comparison of dietary composition of control chow and High Fructose Diet

**Supplementary table 2:** Maternal weight, insulin and cortisol plasma level. Data are expressed as mean± SD and analyzed with Student t-test (** p<0.01).

**Supplementary figure 1:**

| Company | Chow Diet<br>Lab Diet | High-Fructose Diet<br>Research Diets |
|---|---|---|
| **Catalog Number** | 5001 | D05111802 |
| **Carbohydrate (% kCal)** | 57 | 70 |
| **Fructose (% kCal)** | 0.3 | 55 |
| **Fat (% kCal)** | 13 | 10 |
| **Protein (% kCal)** | 30 | 20 |
| **kCal/gram** | 3.35 | 3.85 |

**Supplementary table 1**
